# Dysfunctional synaptic competition at dendritic spines in Fragile X syndrome

**DOI:** 10.64898/2026.02.25.708113

**Authors:** Y. Ramiro-Cortés, A.M. Panzarino, M. Royo, K. Shionoya, I. Israely

**Affiliations:** Champalimaud Research, Champalimaud Centre for the Unknown, 1400-038 Lisbon, Portugal; Department of Pathology and Cell Biology, Department of Neuroscience, in the Taub Institute for Research on Alzheimer’s Disease and the Aging Brain, Columbia University Medical Center, Vagelos College of Physicians and Surgeons, Columbia University, New York, NY 10032, USA; Instituto de Fisiología Celular, Universidad Nacional Autónoma de México, 04510, Ciudad de México, México; Department of Neurobiology and Biophysics, University of Washington, Seattle, WA 98195, USA

## Abstract

Dendritic spines are highly dynamic structures whose morphology and lifespan are modified in response to synaptic efficacy changes. Structural modifications following activity support the long-term encoding of information and could allow for the remodeling of neural circuits. Long-term depression (LTD) is a key mechanism for synaptic weight regulation, yet its structural correlates — particularly for long-lasting, protein synthesis dependent forms — remain poorly understood. Furthermore, in humans, this type of plasticity is often disrupted in neurodevelopmental disorders, correlating with cognitive dysfunction and structural abnormalities. Fragile X Syndrome (FXS) is the most common inherited form of intellectual disability and is characterized by excessive metabotropic receptor-mediated synaptic depression, excessive protein synthesis, and spine abnormalities. Here, we investigate the relationship between long lasting synaptic depression and structural plasticity, as well as the role of protein availability in determining how many spines can simultaneously undergo structural changes during LTD in both healthy and FXS mutant neurons. Using high resolution optical methods, we developed and tested a method for inducing metabotropic glutamate receptor (mGluR)–dependent depression at single spines via glutamate uncaging in mouse hippocampal neurons. We found that this form of activity leads to robust spine shrinkage, which requires new protein synthesis. However, when we induced this depression at multiple spines, they competed for structural changes and only one spine shrank. We hypothesized that this was due to limited resources, in the form of newly made proteins, and therefore, we decided to test if spine competition would be altered in the mouse model of FXS, where protein levels are abnormally elevated. Indeed, we found that competition was absent in FXS mutant neurons, and all of the stimulated spines underwent shrinkage following LTD induction. Importantly, we found that single spine structural plasticity in FXS was expressed to the same degree as in WT controls. Taken together, these findings suggest that the hallmark phenotype of excess mGluR LTD in FXS may result from a greater number of inputs undergoing synaptic depression, rather than excessive LTD at individual synapses. By probing plasticity at the level of individual inputs, our findings highlight the importance of evaluating activity across groups of synapses, in order to uncover plasticity interactions that are critical for learning. Understanding how these mechanisms are disrupted in neurodevelopmental disorders such as FXS can inform the development of effective therapeutic strategies.

## Introduction

Dendritic spines are the principal sites of excitatory synaptic transmission and are highly dynamic structures whose number, size, and shape are modified by neural activity. Spine morphology is closely linked to synaptic function, reflecting correlations between spine size, dendritic location, and synaptic integration (Häusser and Mel, 2003; Magee, 2000; Matsuzaki et al., 2001; Harvey and Svoboda, 2007; Zito et al., 2009). At individual synapses, long-term potentiation (LTP) is robustly accompanied by spine enlargement and stabilization, particularly during the initial expression of potentiation (Matsuzaki et al., 2004), providing a well defined structural correlate, and visual proxy, of synaptic strengthening. By contrast, the structural expression of long-term depression (LTD) remains far less defined.

Although spine shrinkage has been observed following LTD under specific induction paradigms, the conditions that support its persistence and coordination across synapses remain unresolved. Given the prominent role of LTD in shaping synaptic and circuit function, and its dysregulation in multiple neurodevelopmental and neuropsychiatric disorders (Costa-Mattioli and Monteggia, 2013), defining the structural rules governing bidirectional plasticity at spines represents a major unresolved problem.

Structural plasticity unfolds over multiple timescales, and a key organizing principle is its dependence on new protein synthesis. Rapid spine morphological remodeling can occur independently of translation, whereas stabilization of long-lasting structural change, most clearly established for spine growth during synaptic potentiation, requires new protein synthesis (Govindarajan et al., 2011). This protein dependence introduces intrinsic constraints at the level of dendritic domains and was anticipated by models such as synaptic tagging and capture, which propose that synapses compete for access to newly synthesized plasticity-related resources (Frey and Morris, 1997; Govindarajan et al., 2011; Redondo and Morris, 2011; Ramiro-Cortés, Hobbiss and Israely, 2013). Consistent with this framework, synapses have been shown to compete or cooperate for a limited pool of proteins in order to undergo stable long-term structural modifications (Govindarajan et al., 2011). Whether analogous protein-dependent and competitive mechanisms govern the structural expression of synaptic depression is largely unknown.

Group I metabotropic glutamate receptors (Group I mGluRs) provide a powerful model for addressing these questions. Spine shrinkage can be induced by specific patterns of activity, and those that are mediated by NMDA receptor activation and do not recruit new protein synthesis at single spines, may reflect short-lived morphological fluctuations rather than stable remodeling (Holbro et al., 2009; Oh et al., 2012; Hayama et al., 2013). Group I mGluRs, on the other hand, drive a form of synaptic depression that is both long-lasting and protein synthesis dependent, and can be engaged by paired-pulse low-frequency stimulation of hippocampal neurons (Huber et al., 2000; Faas et al., 2002; Nosyreva and Huber, 2005). These changes in efficacy are accompanied by robust spine shrinkage and elimination when pharmacological activation of Group I mGluRs is induced globally (Ramiro-Cortés and Israely, 2013). Despite extensive study at the population level, the structural consequences of functional mGluR-LTD at single spines, and the extent to which they are stabilized by protein synthesis, are yet to be determined. As multiple inputs are typically co-active in vivo, determining whether mGluR-LTD produces persistent, input specific spine shrinkage and how such changes are regulated when activity occurs across neighboring synapses is an open question.

Within a given branch, synapses share access to locally synthesized proteins and compete or cooperate for the stabilization of long-lasting structural changes that are driven by potentiation (Govindarajan et al., 2011). Such interactions may enhance information storage capacity while imposing local limits on plasticity expression. Whether the expression of synaptic depression between neighboring spines is bound by similar interaction rules, particularly in the context of protein synthesis-dependent LTD, is unknown.

Dysregulation of mGluR-dependent LTD and protein synthesis is a recurring feature of neurodevelopmental disorders, prominently including Fragile X syndrome (FXS) (Bear et al., 2004; Bagni and Zukin, 2019; Santini et al., 2014; Costa-Mattioli and Monteggia, 2013). Loss of FMRP, an RNA-binding protein and translational repressor encoded by the *FMR1* gene, leads to elevated protein synthesis, abnormal spine morphology, and exaggerated mGluR-dependent plasticity (Comery et al., 1997; Nimchinsky et al., 2001; Dolen et al., 2007; Jacquemont et al., 2018). In the cortex of the Fmr1 KO mouse, glutamate uncaging revealed stronger synapses resulting from excess connectivity onto single inputs without alterations in spine morphology (Booker et al., 2019). These observations led us to hypothesize that dysregulated protein availability disrupts the normal constraints governing structural plasticity and synaptic competition.

Here, we use single-spine resolution to define the structural correlates of synaptic depression and how these changes are coordinated across synapses. Using an optical paired-pulse low-frequency glutamate uncaging paradigm, we induce mGluR-dependent LTD at single, identified spines. We show that synaptic depression is accompanied by robust, long-lasting spine shrinkage that requires Group I mGluR activation and new protein synthesis. We further demonstrate that when multiple spines are co-activated, they compete for the ability to undergo sustained shrinkage, revealing a structural signature of resource-limited plasticity. Strikingly, this competitive interaction is abolished in a mouse model of Fragile X syndrome, linking dysregulated protein synthesis to impaired synaptic competition. Together, our findings establish dendritic spines as dynamic and competitive units of structural plasticity during synaptic depression and identify a mechanism by which altered protein synthesis disrupts circuit remodeling.

## Results

### Single synapses undergo functional depression and spine shrinkage following optical induction of mGluR-dependent plasticity

To examine the structural correlates of synaptic depression at single inputs, we developed an optical stimulation paradigm that takes advantage of the focal release of the neurotransmitter glutamate through the delivery of paired pulses of light, designed to mimic the electrical paired-pulse low-frequency stimulation known to induce mGluR-dependent LTD (Huber et al., 2000; Kemp and Bashir, 1999). Briefly, upon bath circulation of MNI-glutamate (“caged” glutamate) (2.5mM in ACSF containing 1mM Mg and 2mM Ca; see methods for details), we delivered 90 paired pulses of light to release the active neurotransmitter over 15 minutes at low-frequency (pp-LFU) (50 ms inter-pulse interval, 0.1Hz) to individual dendritic spines on CA1 pyramidal neurons in mouse hippocampal organotypic slice cultures (Fig. 1a). To assess whether this pp-LFU protocol induced synaptic depression, we performed whole-cell patch-clamp recordings at the soma of neurons filled with a fluorescent dye (Alexa 594) and monitored uncaging-evoked EPSCs under voltage-clamp conditions. Following baseline measurements of amplitude obtained by delivering test pulses of glutamate uncaging to multiple spines, we then applied our pp-LFU stimulation to a single target spine (stimulated spine). Subsequent test pulses were then delivered to the stimulated spine and to neighboring unstimulated spines. Robust synaptic depression was observed specifically at the stimulated spine and persisted for the duration of the recording (∼1 h; 54.91 ± 3.26%) while responses at neighboring unstimulated spines remained stable (115.91 ± 4.31%) (Fig. 1b,c). Depression developed rapidly and reached a maximum value of 50.00 ± 8.78% approximately 25 min after stimulation (Fig. 1b,d). All of the experiments were performed in ACSF containing 2 mM CaCl_2_ and 1 mM MgCl_2_.

**Figure 1.**
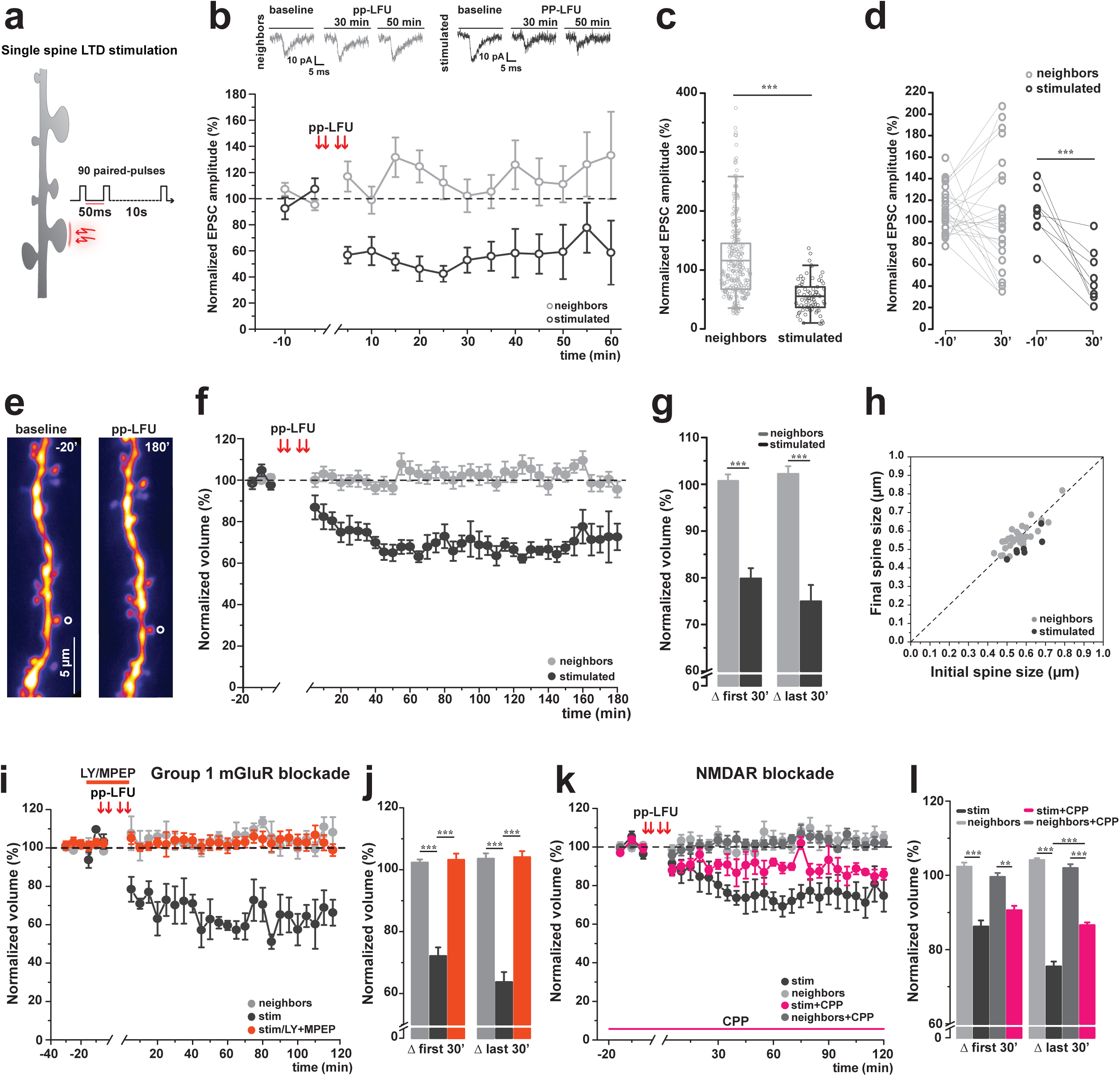
Single-spine mGluR-dependent LTD induces persistent synaptic depression and structural shrinkage. This paradigm produces long-lasting reductions in uEPSC amplitude and spine volume. These changes require Group I mGluR activation and are independent of NMDAR activation. **a)** Schematic of the optically delivered single-spine mGluR-LTD protocol. Two-photon glutamate uncaging delivers paired-pulse low-frequency stimulation (pp-LFU: 90 paired pulses, 50 ms interpulse interval, 0.1 Hz, 15 min) to an individual spine on a CA1 pyramidal neuron in organotypic hippocampal slice culture. **b)** Whole-cell voltage-clamp recordings from CA1 pyramidal neurons in slice cultures from WT C57BL/6J mice. Patched cells were filled with Alexa 594, and uEPSCs were evoked by two-photon glutamate uncaging using 1 ms test pulses. Upper panels: representative uEPSC traces from neighbors (left, grey) and stimulated (right, black) spines before and after pp-LFU induction. Time course of normalized uEPSC amplitude for stimulated spines (black open circles, n=8) and neighboring spines (gray open circles, n=27) from 8 cells. Time points are plotted in 5 min bins. Data represent mean ± SEM. **c)** Normalized uEPSC amplitude post-stimulation for neighboring and stimulated spines. Box plots represent median ± IQR. Kruskal-Wallis test, ***p < 0.0001. **d)** Paired comparison of normalized uEPSC amplitude at baseline (−10 min) and 30 min post-stimulation for individual stimulated (black) and neighboring (gray) spines. Lines connect paired measurements. Paired t-test, ***p = 0.0003. **e)** Representative two-photon z-stack projections of a dendrite from a GFP transfected CA1 neuron during baseline (−20 min) and 180 min post-pp-LFU in hippocampal slice cultures from WT C57BL/6J mice. Stimulated spine indicated by open circle. Scale bar: 5 μm. **f)** Time course of normalized spine volume following pp-LFU for stimulated spines (black filled circles, n=10) and neighboring spines (gray filled circles, n=57) from 10 cells in WT C57BL/6J slice cultures. Time points are plotted in 5 min bins. Data represent mean ± SEM. **g)** Quantification of normalized spine volume shown in (f) averaged over the first 30 min and last 30 min post-pp-LFU for stimulated (black) and neighboring (gray) spines. Bars represent mean ± SEM. One-way ANOVA with Tukey *post hoc*, ***p < 0.0001. **h)** Scatter plot of initial spine size (μm) versus final spine size (180 min) for stimulated spines (black circles, n=10) and neighboring spines (gray circles, n=57). Dashed line indicates no change. **i)** Time course of normalized spine volume following pp-LFU under Group I mGluR blockade with LY367385 (100 μM) and MPEP (10 μM). Stimulated spines (orange circles, n=4) and neighboring spines (gray circles, n=24) from 4 cells. Interleaved control stimulated spines without blockade (black circles, n=3 spines, 3 cells). Data represent mean ± SEM. **j)** Quantification of normalized spine volume averaged over the first 30 min and last 30 min post-pp-LFU for conditions shown in (i). Bars represent mean ± SEM. One-way ANOVA with Tukey *post hoc*, ***p < 0.0001. **k)** Time course of normalized spine volumes following pp-LFU in the presence of the NMDAR antagonist CPP (5 μM). Stimulated spines (magenta circles, n=3) and neighboring spines (dark grey circles, n=19) from 3 cells. Interleaved control stimulated spines with no drug (black circles, n=3) and neighboring spines (light gray circles, n=16) from 3 cells. Data represent mean ± SEM. **l)** Quantification of normalized spine volumes averaged over the first 30 min and last 30 min post-pp-LFU for all conditions shown in (k). Bars represent mean ± SEM. One-way ANOVA with Tukey *post hoc*, **p < 0.001, ***p < 0.0001.

Having successfully demonstrated that individual synapses functionally depress upon pp-LFU stimulation, we next examined the structural correlates of this plasticity over time by tracking spine volume changes after glutamate uncaging induced LTD (Fig. 1e). Two-photon imaging was used to record spine volumes first during a baseline period and then following the induction of pp-LFU at a select spine and its unstimulated neighbors for up to 3 hours. We found that stimulated spines exhibited robust and persistent shrinkage (74.95 ± 15.98%), whereas neighboring spine volumes remained stable (102.23 ± 14.44%) (Fig. 1f,g). Spine shrinkage was observed across various initial sizes (Fig. 1h).

To test whether pp-LFU induced plasticity depends on activation of Gp1 mGluRs, we performed stimulation in the presence of the selective mGluR1 and mGluR5 antagonists LY367385 and MPEP, respectively. Pharmacological blockade of Gp1 mGluRs during pp-LFU completely abolished spine shrinkage (103.20 ± 2.25%). In the absence of these antagonists, pp-LFU led to robust spine shrinkage (63.75 ± 3.18% of stimulated spines) (Fig. 1i,j), establishing that Group I mGluR signaling is required. However, since glutamate uncaging activates multiple glutamate receptor classes at the stimulated spine, albeit with distinct activation kinetics, we next asked whether NMDA receptors also contribute. NMDAR blockade with CPP significantly attenuated, but did not eliminate, spine shrinkage (86.61 ± 0.07% compared to 75.51 ± 1.30% without blockade) (Fig. 1k,l). These data demonstrate that pp-LFU induced spine shrinkage is strictly dependent on Group I mGluR activation, while NMDAR co-activation enhances the induction of structural plasticity, but is not necessary for its expression. This suggests a synergistic interaction between mGluR and NMDAR signaling at the stimulated spine, although mGluR activation is required to drive persistent spine shrinkage.

### Spine shrinkage during mGluR activation requires protein synthesis

A defining feature of metabotropic glutamate receptor-dependent synaptic plasticity is its reliance on new protein synthesis to support long-lasting changes in synaptic efficacy (Huber KM et al., 2000). To test whether protein synthesis is required for synaptic depression and the associated spine shrinkage at individual synapses, we performed pp-LFU experiments under conditions of translational blockade. Hippocampal slice cultures were pre-incubated with the protein synthesis inhibitor anisomycin for 20 min prior to induction of plasticity. Under these conditions, pp-LFU failed to induce synaptic depression at stimulated spines, as measured by uEPSC amplitudes before and after stimulation (95.09 ± 2.4% at stimulated spines or 94.84 ± 3.05% at neighbors) (Fig. 2b–d). Thus, protein synthesis is required for pp-LFU induced functional depression at single synapses.

**Figure 2.**
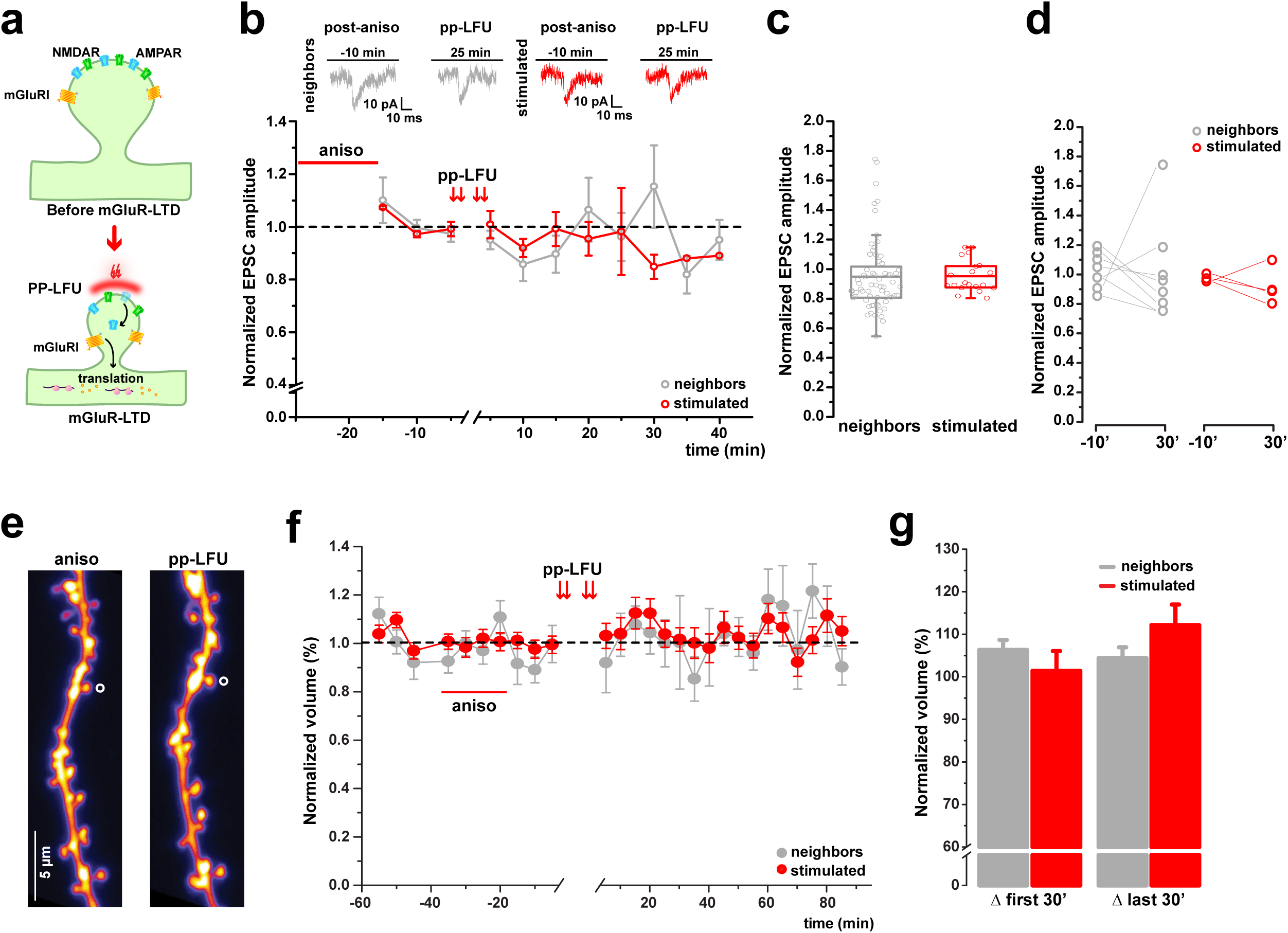
De novo protein synthesis is required for mGluR-dependent LTD and long-lasting spine shrinkage. Inhibition of translation prevents sustained synaptic depression and spine shrinkage following mGluR-dependent LTD. **a)** Schematic of mGluR-dependent signaling leading to local protein synthesis during LTD induced by pp-LFU at a single spine. **b)** Whole-cell voltage-clamp recordings from CA1 pyramidal neurons in slice cultures from WT C57BL/6J mice following pre-incubation with the protein synthesis inhibitor anisomycin (50 μM, 20 min pre-incubation). Upper panels: representative uEPSC traces from neighbors (left, gray) and stimulated (right, red) spines before and after pp-LFU. Time course of normalized uEPSC amplitude for stimulated spines (red circles, n=4) and neighboring spines (gray circles, n=13) from 4 cells. Data represent mean ± SEM. **c)** Normalized uEPSC amplitude post-stimulation for neighboring and stimulated spines under anisomycin treatment. Box plots represent median ± IQR. Kruskal-Wallis test, p=0.275. **d)** Paired comparison of normalized uEPSC amplitude at baseline (−10 min) and 30 min post-stimulation for individual stimulated (red) and neighboring (gray) spines. Lines connect paired measurements. Paired t-test, p=0.17. **e)** Representative two-photon z-stack projections of a dendrite from a GFP-transfected CA1 neuron during baseline (-20 min) and 120 min post-pp-LFU under anisomycin treatment in WT C57BL/6J hippocampal slice cultures. Stimulated spine indicated by open circle. Scale bar: 5 μm. **f)** Time course of normalized spine volume following pp-LFU under anisomycin blockade (50 μM, 20 min pre-incubation). Stimulated spines (red circles, n=6) and neighboring spines (gray circles, n=38) from 6 cells in hippocampal slice cultures from WT C57BL/6J mice. Data represent mean ± SEM. **g)** Quantification of normalized spine volume averaged over the first 30 min and last 30 min post-pp-LFU for stimulated (red) and neighboring (gray) spines. Bars represent mean ± SEM. One-way ANOVA with Tukey *post hoc*, p=0.835.

We next examined the structural consequences of pp-LFU during translational blockade using two-photon imaging over the course of several hours (Fig. 2e–g). In contrast to control conditions, when anisomycin was applied, pp-LFU failed to induce spine shrinkage at stimulated inputs (105.03 ± 4.18%) and volumes were similar to those of neighboring unstimulated spines (102.61 ± 2.06%) (Fig. 2f,g). Together, these results demonstrate that protein synthesis is required for both synaptic depression and spine shrinkage induced by pp-LFU.

### Competition between co-active spines during synaptic depression

We showed that induction of metabotropic glutamate receptor driven synaptic depression at a single input leads to robust functional depression accompanied by spine shrinkage. It remained unclear, however, whether similar structural outcomes would be expressed when multiple synaptic inputs are activated concurrently. Previous work demonstrated that during protein synthesis dependent LTP, neighboring spines compete for growth when multiple inputs are co-active (Govindarajan et al., 2011). To test whether structural depression is similarly constrained, we designed an experiment in which two neighboring spines were stimulated pseudo-synchronously using the pp-LFU paradigm. Paired pulses were delivered in an interleaved manner between two spines on the same dendritic branch, such that each spine received 90 paired pulses of glutamate uncaging, offset by 50 ms, over a total stimulation period of 15 min (Fig. 3a). Stimulated spines were located approximately 20 µm apart on the same secondary or tertiary dendrites, and structural changes were monitored for up to 2 h following stimulation (Fig. 3a–d).

**Figure 3.**
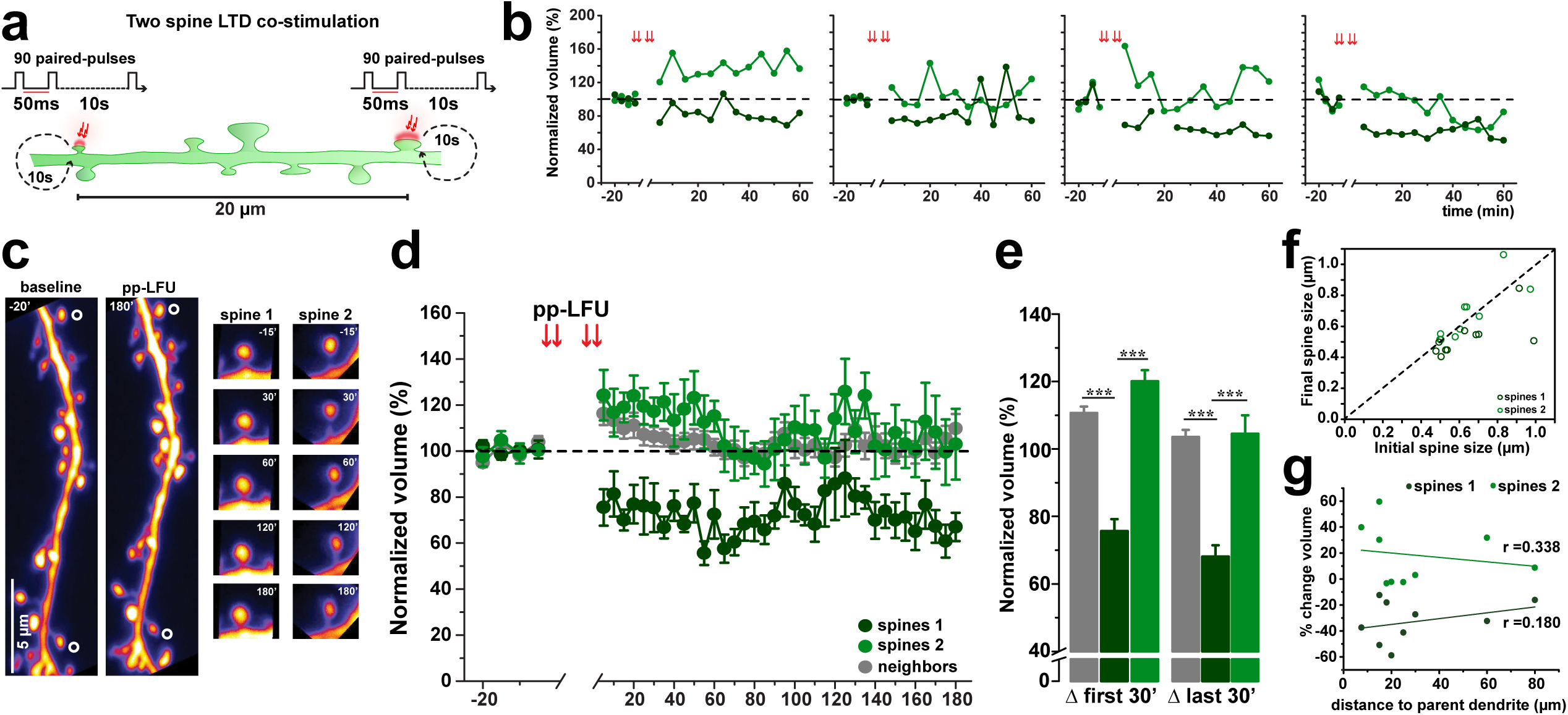
Simultaneous LTD induction at multiple spines triggers structural competition. Co-stimulated spines display asymmetric long-term structural changes following LTD induction. **a)** Schematic of the two-spine competition experiment. Two spines on the same dendrite, separated by ∼20 μm, are co-stimulated with interleaved pp-LFU. One spine receives a pair of pulses, followed 100 ms later by a pair of pulses to the second spine, repeating at 0.1 Hz for 15 min. Each spine receives 90 pairs (180 total pulses). **b)** Individual examples of two-spine competition experiments showing normalized spine volume changes in both spines over the first hour post-pp-LFU. Each panel represents one experiment. Light and dark green traces represent the two co-stimulated spines, group data in (d). **c)** Representative two-photon z-stack projections of a dendrite from a GFP-transfected CA1 neuron during baseline (−20 min) and 180 min post-pp-LFU in WT C57BL/6J hippocampal slice cultures. Co-stimulated spines indicated by open circles. Right panels: time-lapse of individual spine 1 and spine 2 at selected time points up to 3 hours post stimulation. Scale bar: 5 μm. **d)** Time course of normalized spine volume following two-spine pp-LFU. Spines were classified *post hoc* as Spine 1 (dark green, exhibiting ≥20% shrinkage within the first 15 min that was maintained for the duration of the experiment) or Spine 2 (light green, not meeting this criterion). Neighboring spines shown in gray. n=10 cells, 20 stimulated spines (10 Spine 1, 10 Spine 2), 47 neighboring spines. Data represent mean ± SEM. **e)** Quantification of normalized spine volume averaged over the first 30 min and last 30 min post-pp-LFU for Spine 1 (dark green), Spine 2 (light green), and neighboring spines (gray). Bars represent mean ± SEM. One-way ANOVA with Tukey *post hoc*, ***p < 0.0001. **f)** Scatter plot of initial spine size (μm) versus final spine size (180 min) for Spine 1 (dark green) and Spine 2 (light green). Dashed line indicates no change. **g)** Percentage volume change as a function of distance to parent dendrite branch point (μm) for Spine 1 (dark green, r=0.338) and Spine 2 (light green, r=0.180). Linear fits shown.

Whereas single spine stimulation reliably induced spine shrinkage, simultaneous stimulation of two spines produced asymmetric structural outcomes (Fig. 3b,d). In each experiment, one stimulated spine underwent rapid and sustained shrinkage, while the other showed variable responses ranging from transient growth to volume fluctuations (Fig. 3b). Based on the structural response, each spine was classified *post hoc*: a spine that exhibited shrinkage within the first 15 minutes that persisted for the duration of the recording was designated Spine 1, while a spine that did not show early shrinkage was designated Spine 2 (Fig. 3e). The Spine 1 group decreased in volume rapidly and remained persistently smaller (Spine 1 group: 75.63 ± 1.95%, first 30 min; 68.09 ± 3.34%, last 30 min). In contrast, the Spine 2 group exhibited a modest, transient increase in volume after stimulation that gradually declined (Spine 2 group: 120.63 ± 3.33%, first 30 min; 104.51 ± 5.48%, last 30 min). In a few cases, a spine initially classified as Spine 2 subsequently exhibited delayed shrinkage (Fig. 3b), although the predominant outcome remained asymmetric plasticity.

We next examined whether anatomical or stimulation-related factors predicted which spine underwent shrinkage. No correlation was observed between a spine’s distance from the parent dendrite and its structural outcome (Spine 1: r = 0.180; Spine 2: r = 0.338; Fig. 3g). The likelihood of a spine undergoing shrinkage was not correlated with its initial volume (Fig. 3f). Together, these data show that when two synaptic inputs undergo pp-LFU–induced LTD simultaneously, structural depression is expressed asymmetrically, consistent with competition for limited plasticity resources within a dendritic compartment.

### Dysfunctional synaptic competition in Fragile X syndrome

The competitive interactions observed between co-active spines during synaptic depression point to a constraint that limits how many inputs can express long-lasting plasticity concurrently. Given the requirement for new protein synthesis during mGluR-dependent LTD, we hypothesized that increasing the availability of translational products might alleviate this constraint. To test this idea, we turned to a mouse model of Fragile X syndrome, in which loss of the RNA-binding protein FMRP results in elevated basal protein synthesis (Osterweil et al., 2010). We generated hippocampal organotypic slice cultures from Fmr1 knockout mice (Fmr1 KO) and wild-type littermates (Fmr1 WT) and examined pp-LFU induced structural plasticity at pairs of neighboring spines.

### Loss of competition between co-active spines in Fmr1-KO neurons

When two neighboring spines in Fmr1 KO neurons were stimulated pseudo-synchronously using the pp-LFU paradigm (Fig. 4a), both spines reliably underwent shrinkage – in striking contrast to the asymmetric outcomes observed in wild-type neurons (Fig. 4f–i). Spine volumes decreased within the first 30 min after stimulation (Spine 1: 85.06 ± 3.22%; Spine 2: 85.36 ± 3.36%) and stabilized over the subsequent 2 h at levels comparable to single-spine stimulation (Spine 1: 64.58 ± 2.06%; Spine 2: 70.03 ± 1.70%). Thus, co-active spines in Fmr1 KO neurons expressed structural depression concurrently, indicating a loss of the competitive constraint that limits plasticity in wild-type neurons.

**Figure 4.**
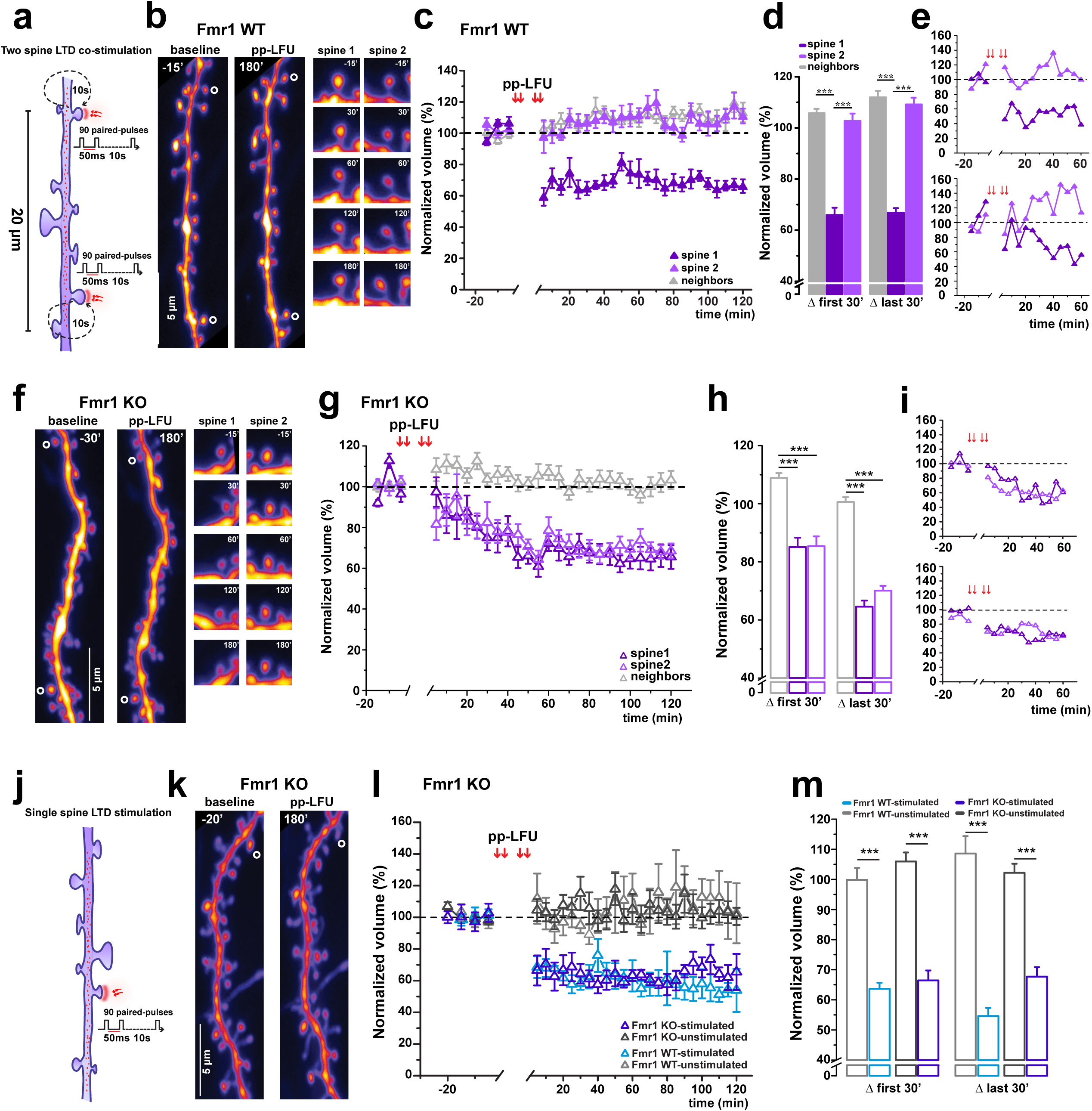
Fragile X disrupts structural competition between co-stimulated spines. In Fmr1 KO neurons, both stimulated spines exhibit persistent structural shrinkage following LTD induction in contrast to the asymmetric response observed in WT neurons. **a)** Schematic of the two-spine competition experiment in Fmr1 WT and Fmr1 KO hippocampal slice cultures. Protocol as described in Figure 3a. **b)** Representative two-photon z-stack projections of a dendrite from a GFP-transfected CA1 neuron in Fmr1 WT slice cultures during baseline (−15 min) and 180 min post-pp-LFU. Co-stimulated spines indicated by open circles. Right panels: time-lapse of individual Spine 1 and Spine 2 at selected time points. Scale bar: 5 μm. **c)** Time course of normalized spine volume following two-spine pp-LFU in Fmr1 WT neurons. Spine 1 (dark purple, filled triangles), Spine 2 (light purple, filled triangles), and neighboring spines (gray, open triangles). n=8 cells, 16 stimulated spines (8 Spine 1, 8 Spine 2), 38 neighboring spines. Data represent mean ± SEM. **d)** Quantification of normalized spine volume averaged over the first 30 min and last 30 min post-pp-LFU for Spine 1 (dark purple), Spine 2 (light purple), and neighboring spines (gray) in Fmr1 WT. Bars represent mean ± SEM. One-way ANOVA with Tukey *post hoc*, ***p < 0.0001. **e)** Individual examples of two-spine competition experiments in Fmr1 WT showing normalized spine volume changes in both spines over the first 60 min post-pp-LFU. Group data in (c). **f)** Representative two-photon z-stack projections of a dendrite from a GFP-transfected CA1 neuron in Fmr1 KO slice cultures during baseline (−30 min) and 180 min post-pp-LFU. Co-stimulated spines indicated by open circles. Right panels: time-lapse of individual Spine 1 and Spine 2. Scale bar: 5 μm. **g)** Time course of normalized spine volume following two-spine pp-LFU in Fmr1 KO neurons. Spine 1 (dark purple, open triangles), Spine 2 (light purple, open triangles), and neighboring spines (gray, open triangles). n=13 cells, 26 stimulated spines (13 Spine 1, 13 Spine 2), 27 neighboring spines. Data represent mean ± SEM. **h)** Quantification of normalized spine volume averaged over the first 30 min and last 30 min post-pp-LFU for Spine 1 (dark purple, open), Spine 2 (light purple, open), and neighboring spines (gray, open) in Fmr1 KO. Bars represent mean ± SEM. One-way ANOVA with Tukey *post hoc*, ***p < 0.0001. **i)** Individual examples of two-spine competition experiments in Fmr1 KO showing normalized spine volume changes in both spines over the first 60 min post-pp-LFU. Group data in (g). **j)** Schematic of single-spine pp-LFU stimulation in Fmr1 KO hippocampal slice cultures. Protocol as described in Figure 1a. **k)** Representative two-photon z-stack projections of a dendrite from a GFP-transfected CA1 neuron in Fmr1 KO slice cultures during baseline (−20 min) and 180 min post-pp-LFU. Stimulated spine indicated by open circle. Scale bar: 5 μm. **l)** Time course of normalized spine volume changes following the stimulation of a single spine to undergo depression by pp-LFU in either Fmr1 KO neurons or in control Fmr1 WT littermate neurons. Stimulated spines in Fmr1 KO (dark purple triangles, n=5) and neighboring spines (dark gray triangles, n=24) from 5 cells. Stimulated spines from Fmr1 WT (light blue triangles, n=3) and neighboring spines (light gray triangles, n=12) from 3 cells. Data represent mean ± SEM. **m)** Quantification of normalized spine volume averaged over the first 30 min and last 30 min after pp-LFU for stimulated (Fmr1 KO: dark purple, Fmr1 WT: light blue) and unstimulated spines (Fmr1 KO: dark gray, Fmr1 WT: light gray). Bars represent mean ± SEM. One-way ANOVA with Tukey *post hoc*, ***p < 0.0001.

All experiments were conducted blind to genotype and analyzed prior to group assignment. Spines were classified *post hoc* using the same criteria across genotypes: Spine 1 (exhibiting early and sustained shrinkage) and Spine 2 (not meeting this criterion; see Methods). Following unblinding, we confirmed that Fmr1 WT neurons exhibited competitive structural outcomes indistinguishable from those observed in C57BL/6J wild-type neurons. When two spines were co-stimulated in Fmr1 WT neurons, one spine consistently underwent sustained shrinkage (Spine 1) while the other remained near baseline (Spine 2) (Fig. 4b-e). Spine 1 decreased in volume during the first 30 min (65.95 ± 2.79%) and remained depressed at later time points (66.84 ± 1.80%), while Spine 2 showed no significant change (102.74 ± 2.87% first 30 min; 109.16 ± 2.64% last 30 min). These results rule out genetic background effects and confirm that loss of competition is specific to the Fmr1 null condition.

### Single-spine structural plasticity is preserved in Fmr1-KO neurons

To determine whether the absence of competition in Fmr1 KO neurons reflected a general alteration in spine plasticity, we next examined structural changes induced by pp-LFU at single spines (Fig. 4j). Individual spines in both Fmr1 KO and Fmr1-WT neurons underwent robust and sustained shrinkage following stimulation (Fig. 4k-m). The magnitude and time course of spine shrinkage were comparable between genotypes (Fmr1 KO: 66.46 ± 3.31% early, 67.72 ± 3.16% late; Fmr1 WT: 63.74 ± 2.05% early, 54.64 ± 2.69% late), while unstimulated neighboring spines did not shrink in either case. Together with the finding that competition is lost in Fmr1 KO spines, these data indicate that the excess mGluR-LTD observed at the population level in FXS does not reflect exaggerated plasticity at individual synapses, but rather a failure to constrain how many synapses undergo depression simultaneously.

## Discussion

Synaptic plasticity is expressed at individual dendritic spines, yet learning emerges from interactions among many synapses within dendritic compartments. While long term potentiation and depression are well established mechanisms for modifying synaptic strength, far less is known about how plasticity at one spine influences the expression of plasticity at neighboring inputs, or how such interactions are limited by local molecular resources. Understanding these inter-spine learning rules is particularly important for forms of plasticity that require new protein synthesis, where the availability of translational resources may limit or bias which synapses undergo lasting modification.

Although synaptic plasticity has often been framed in terms of LTP, substantial evidence supports LTD as an equally important mechanism for learning and memory (Bear, 1996; Malenka and Bear, 2004). Here, we establish a single-spine optical stimulation paradigm that enables synaptic depression to be induced and monitored with sufficient resolution to examine inter-spine interactions directly. Using paired-pulse low-frequency glutamate uncaging (pp-LFU), we demonstrate that individual dendritic spines undergo persistent functional depression together with stable spine shrinkage under physiological conditions, without pharmacological manipulation of NMDA receptors. This form of plasticity is protein synthesis dependent and requires Group I mGluR activation, with a modulatory contribution from NMDA receptors. By resolving plasticity at the level of individual inputs, this approach reveals competitive interactions between neighboring spines that shape the expression of LTD within dendritic compartments and are disrupted in a Fragile X syndrome model.

### Protein synthesis dependent mGluR-LTD at single synapses

New protein synthesis is critical both for the establishment of long lasting plasticity and for regulating its expression during potentiation (Frey and Morris, 1997; Klann and Dever, 2004; Govindarajan and Israely, 2010), leading us to focus on mGluR mediated LTD, a form of depression known to require translation (Huber et al., 2000; Nosyreva and Huber, 2005). Our previous work showed that global induction of mGluR-dependent depression drives spine shrinkage, but it remained unclear whether such structural plasticity could be induced at individual synaptic inputs or required coordinated activation of multiple synapses (Ramiro-Cortés and Israely, 2013). Prior single spine LTD studies relied on NMDA receptor activation through depolarization or back-propagating action potentials (Oh et al., 2013; Hayama et al., 2013), or pharmacologically isolated mGluR-dependent depression without assessing structural consequences (Holbro et al., 2009). Our optical pp-LFU paradigm resolves this limitation: it induces correlated functional depression and persistent spine shrinkage at individual spines, both of which require new protein synthesis. This establishes spine volume as a reliable proxy for synaptic weakening at single inputs, analogous to optical readouts of potentiation.

Pharmacological dissection revealed that Group I mGluR activation is required for pp-LFU-induced plasticity, while NMDAR blockade attenuated but did not abolish structural depression. This is consistent with a cooperative signaling model in which mGluR-driven plasticity is modulated by NMDAR activity, possibly through non-ionotropic NMDAR signaling (Nabavi et al., 2013; Stein et al., 2015; Doré et al., 2017). The delayed onset of structural depression under NMDAR blockade suggests that NMDAR co-activation accelerates the induction of spine shrinkage but is not required, as mGluR signaling alone is sufficient given adequate time. By maintaining the native signaling environment rather than pharmacologically isolating receptor classes, our paradigm preserves the convergence of mGluR and NMDAR signaling at the stimulated spine, providing a mechanistic basis for integrating synaptic depression across neighboring inputs and allowing competition or coexistence of plasticity to emerge depending on local protein availability rather than engagement of a single receptor pathway.

### Inter-spine competition during protein synthesis dependent LTD

Dendritic spines do not function as isolated units but interact within dendritic compartments, competing or cooperating for access to plasticity related resources. Our previous work established such competitive interactions during protein synthesis dependent LTP, in which co-active neighboring spines within a dendritic branch influence one another’s long-term expression of potentiation (Govindarajan et al., 2011). Whether similar constraints operate during synaptic depression has remained unclear. Here, we show that when two neighboring inputs simultaneously undergo mGluR-dependent LTD, structural outcomes are consistently asymmetric: one spine undergoes shrinkage, whereas the other remains unchanged or slightly increases in volume. This asymmetry is not explained by initial spine size, distance from the parent dendrite, or stimulation order, indicating that competition for limited local resources constrains the expression of LTD, as it does LTP, at the level of individual inputs.

### Fragile X reveals protein availability as a constraint on synaptic competition

To determine how altered translational capacity influences protein availability and competitive dynamics during mGluR-dependent LTD, we examined neurons from a mouse model of Fragile X syndrome. Fragile X loss of function mice exhibit excessive protein synthesis due to loss of the RNA-binding protein FMRP and display enhanced mGluR-dependent LTD (Huber et al., 2002; Bear et al., 2004). We reasoned that elevated protein availability might relieve the normal constraints that enforce competition between neighboring synapses. Consistent with this prediction, when two nearby spines simultaneously underwent pp-LFU induced LTD in Fmr1 KO neurons, both spines reliably shrank, abolishing the competitive asymmetry observed in wild-type neurons. This is despite normal depression at single Fmr1 KO spines relative to controls. These findings explain the exaggerated mGluR-LTD phenotype in Fragile X: rather than reflecting stronger depression at individual inputs, excess LTD arises from a failure to constrain how many synapses undergo depression concurrently. The basis for this lies in the dynamics of local protein synthesis. Newly synthesized proteins are captured by active synapses to stabilize short-lasting plasticity into enduring synaptic modification (Frey and Morris, 1997), yet local translational resources are finite, forcing co-active inputs to compete for their usage (Govindarajan et al., 2011), ultimately influencing which synapses are recruited into a memory trace (Rogerson et al., 2014); in the absence of FMRP, this selectivity is lost, with direct consequences for how plasticity is distributed across synapses within a dendritic compartment.

Notably, under all conditions tested, pp-LFU induced LTD resulted in persistent spine shrinkage without overt spine elimination. This differs from our previous finding that global mGluR activation via DHPG induced both spine shrinkage and elimination (Ramiro-Cortés and Israely, 2013), consistent with the possibility that global receptor activation engages distinct regulatory mechanisms from those operating at individual synapses. The competitive interactions identified in this study do not reflect synapse removal, and instead establish stable differences in synaptic weight among nearby inputs. Such a plasticity regime supports learning through graded reweighting of synapses within dendritic compartments, rather than through circuit pruning. Spine elimination may represent a distinct outcome that emerges under different activity patterns, developmental stages, or longer timescales than those examined here.

At the population level, competition between multiple synaptic pathways was first observed in the context of synaptic tagging and capture (Fonseca et al., 2004). Our work reveals that neighboring spines within a dendritic branch compete for locally synthesized proteins, and that this competition determines which inputs undergo lasting modification, expressed as stable, bidirectional changes in both synaptic strength and spine structure (Govindarajan et al., 2011; Ramiro-Cortés and Israely, 2013; Ramiro-Cortés, Hobbiss, and Israely, 2014). Consistent with this framework, pharmacological evidence has shown that mTORC1-dependent protein synthesis is required for spine structural changes under normal conditions but is dispensable when translation is chronically elevated in FXS (Thomazeau et al., 2020). It is worth noting that structural modifications following global receptor activation may engage distinct regulatory mechanisms from those operating at individual synapses, as we observed with DHPG mediated spine elimination (Ramiro-Cortés and Israely, 2013). Our data provide the functional consequence of dysregulated protein homeostasis at single-spine resolution, demonstrating that excess translation removes the competitive interactions that normally determine which synapses undergo lasting modification.

Our single spine uncaging approach reveals inter-spine interactions that are obscured when plasticity is assessed at the pathway or whole-neuron level and provides a framework for understanding how dysregulated translation perturbs synaptic competition across neurodevelopmental disorders. Indeed, application of our pp-LFU paradigm to a Shank3 model of ASD revealed distinct alterations in spine structural plasticity (Perera-Murcia et al., 2026), suggesting that this approach may be broadly applicable for probing synaptic plasticity rules across neurodevelopmental disorders. Long-term synaptic modification is not determined solely at individual inputs, but emerges through competitive interactions among neighboring spines, governed by the local availability of newly synthesized proteins. Future work dissecting the distinct contributions of mGluR1 and mGluR5 to input-specific plasticity may reveal additional layers of regulation that shape these competitive outcomes.

## Methods

### Organotypic hippocampal slice culture preparation and transfection

Organotypic hippocampal slice cultures were prepared from postnatal day 7-10 C57BL/6J and Fmr1 KO littermate mice (Jackson Laboratory), as previously described (Govindarajan, et al. 2011). Briefly, hippocampal slices (350 μm thick) were cut with a tissue chopper in ice-cold ACSF containing (in mM): 2.5 KCl, 26 NaHCO_3_, 1.15 NaH_2_PO_4_, 11 D-glucose, 24 sucrose, 1 CaCl_2_, and 5 MgCl_2_. Slices were cultured on membrane inserts (PICMORG50, Millipore) in an interface configuration with culture medium containing: 1× MEM (Invitrogen), 20% horse serum (Invitrogen), 1 mM GlutaMAX (Invitrogen), 27 mM D-glucose, 30 mM HEPES, 6 mM NaHCO_3_, 1 mM CaCl_2_, 1 mM MgSO_4_, 1.2% ascorbic acid, and 1 μg/ml insulin. The pH was adjusted to 7.3 and osmolarity to 300–310 mOsm. All chemicals were from Sigma unless otherwise indicated. Cultures were transfected using a Helios gene gun (Bio-Rad) after 5–6 days in vitro (DIV). Gold beads (10 mg, 1.6 μm diameter, Bio-Rad) were coated with 100 μg AFP plasmid DNA,a GFP-expressing plasmid driven by the β-actin promoter (Inouye et al., 1997) and delivered biolistically at 180 psi.

### Two-photon imaging

Two-photon imaging experiments were performed 2–7 days post-transfection. Slices were perfused with ACSF containing (in mM): 127 NaCl, 2.5 KCl, 25 NaHCO_3_, 1.25 Na H_2_PO_4_, 25 D-glucose, 2 CaCl_2_, and 1 MgCl_2_ (and oxygenated with 95% O_2_/5% CO_2_) at room temperature at a rate of 1.5 ml/min. Imaging was performed using a dual-galvanometer scanning system (Prairie Technologies/Bruker) mounted on a BX61WI Olympus microscope with two Ti:sapphire lasers (Chameleon Ultra II, Coherent; and Mai Tai HP, Spectra Physics): one for imaging (910 nm for AFP, 820 nm for Alexa 594) and one for photoactivation of MNI-L-caged glutamate (720 nm), controlled by PrairieView software. Slices were submerged in oxygenated ACSF and perfused for a minimum of 1 h before the start of imaging sessions. Secondary or tertiary apical dendrites of CA1 neurons were imaged using a water immersion objective (60x, 1.0 NA, Olympus LUMPlan FLN) with a digital zoom of 7x. Z-stack projections (0.3 μm step size) were collected every 5 min. For each experiment, a lower-magnification z-stack was also collected in order to measure spine distance from the parent dendrite.

### Caged glutamate preparation and calibration

Uncaging experiments and caged glutamate calibration were carried out as previously described (Govindarajan et al., 2011), and briefly as follows. MNI-caged-L-glutamate (MNI-Glu; Tocris) was reconstituted in the dark in ACSF lacking CaCl_2_ and MgCl_2_ to make a 10 mM stock solution, which was aliquoted into individual vials for single experiments and stored in the dark at −80°C. Individual aliquots were thawed and diluted to a working concentration of 2.5 mM MNI-Glu in ACSF (uACSF, uncaging ACSF) (see below) prior to each experiment. For each working stock batch, one aliquot was used to validate proper dilution and efficacy: five uncaging test pulses (720 nm, 1 ms) were delivered to single spines and evoked uEPSCs were measured by whole-cell patch clamp recordings. The laser power was adjusted (∼30 mW at the back aperture) to produce uEPSCs comparable in magnitude to endogenous miniature EPSCs, as previously described (Govindarajan et al., 2011).

### Uncaging LTD induction

Uncaging ACSF (uACSF) containing: 2.5 mM MNI-Glu in standard ACSF with 2 mM CaCl_2_ and 1 mM MgCl_2_, was recirculated in a 3 ml volume for 5 min prior to stimulation. The uncaging laser was directed 0.5 μm from the center of the spine head.

The pp-LFU protocol was designed based on electrical paired-pulse low-frequency stimulation protocols (Huber et al., 2000). We delivered 90 paired pulses (50 ms interpulse interval, 0.5 ms pulse width) at 0.1 Hz for 15 min (∼15 mW laser power at the back aperture, 720 nm). After stimulation, perfusion was returned to normal ACSF for the remainder of the experiment. Measurements were acquired at discrete intervals and plotted in 5 min bins, with the first post-stimulation point shown at t = 5 min (representing the initial 0–5 min period after stimulation).

For two-spine competition experiments, two spines on the same dendrite, separated by approximately 20 μm and located in the same focal plane, were selected. Paired pulses were delivered in an interleaved manner: one spine received a pair of pulses (two pulses, 50 ms interpulse interval), followed 100 ms later by delivery of a pair of pulses to the other spine, repeating at 0.1 Hz for 15 min. The stimulation laser returned to the first spine 9.9 s after the second spine was stimulated. Each spine received a total of 90 pairs (180 pulses).

### Pharmacological treatments

For experiments involving pharmacological blockade, slices were pre-incubated with the relevant agents prior to stimulation. Group I mGluR antagonists LY367385 (100 μM; mGluR1) and MPEP (10 μM; mGluR5) were bath-applied for 30 min and included in the uACSF. The NMDAR antagonist CPP (5 μM) was bath-applied for 30 min, included in the uACSF, and maintained throughout the experiment. The protein synthesis inhibitor anisomycin (50 μM) was bath-applied for 20 min before pp-LFU to ensure complete translational inhibition prior to stimulation. Anisomycin was then washed out for 15 min before delivery of pp-LFU, as translational inhibition persists well beyond washout (Frey et al., 1988; Kauderer and Kandel, 2000).

### Electrophysiology

For combined electrophysiology and imaging experiments, whole-cell voltage-clamp recordings were obtained from non-transfected CA1 pyramidal neurons in slice cultures. Borosilicate glass electrodes (8–9 MΩ) were filled with internal solution containing (in mM): 135 K-gluconate, 10 HEPES, 10 Na-phosphocreatine, 3 Na-L-ascorbate, 4 MgCl2, 4 Na-ATP, 0.4 Na-GTP, and 0.025 Alexa 594; pH adjusted to 7.2 with NaOH, osmolarity 292 mOsm. After achieving whole-cell configuration, cells were allowed to fill with Alexa 594 for a minimum of 5 min before beginning baseline imaging and uEPSC measurements. Cells were voltage-clamped at −65 mV. Recordings with series resistance exceeding 25 MΩ were discarded, and series resistance stability was monitored throughout (±20% tolerance). Signals were acquired using a Multiclamp 700B amplifier (Molecular Devices) and digitized with a Digidata 1440 at 3 kHz. uEPSC amplitudes were analyzed using custom code in MATLAB. For experiments conducted under protein synthesis blockade, pre-incubation and washout prior to pp-LFU was done following the same experimental design as for imaging described above.

### Data analysis and Statistics

Spine volumes were measured using SpineS software (Argunşah et al., 2022), which performs automated segmentation of individual spines to extract volume measurements from z-stack projections. Distance from each stimulated spine to the parent dendrite branch point was measured from lower-magnification z-stack projections using PrairieView software.

All spine volume and uEPSC measurements were normalized to the average of baseline values for each spine. For time-course plots, data are presented as mean ± SEM. For quantification of plasticity magnitude, the first 30 min and last 30 min post-stimulation were averaged for each spine and compared across conditions.

In two-spine competition experiments, spines were classified *post hoc* as Spine 1 (exhibiting at least 20% reduction from baseline volume within the first 15 min post-stimulation that was maintained for the duration of the experiment) or Spine 2 (not meeting this criterion). When both spines met the criterion, as was typically the case in Fmr1 KO neurons, designation was assigned by stimulation order.

Statistical comparisons were performed using one-way ANOVA with Tukey *post hoc* test for normally distributed data, Kruskal-Wallis test for non-normally distributed data, and paired t-test for within-spine comparisons of baseline versus post-stimulation values. Normality was assessed using Shapiro-Wilk test. A significance threshold of p ≤ 0.05 was applied. Sample sizes are reported as number of spines, cells, and independent slice cultures in figure legends.

Blinding: Fmr1 KO and wild-type littermate pups were genotyped by tail biopsy prior to slice culture preparation. Pups were coded by litter and number, and genotype information was withheld from the experimenter. Slices from each animal were maintained on separate culture inserts, and experiments were interleaved across genotypes. Unblinding was only done after data analysis was complete.

All animal procedures were performed in accordance with institutional guidelines and approved by the Institutional Animal Care and Use Committees (or equivalent) at the Champalimaud Foundation, Columbia University, and the University of Washington.

## Acknowledgments

We thank Ali Ozgur Argunsah for advice and assistance with data analysis. We also thank Rui Costa, Astra Bryant, and members of the Israely laboratory for their support, helpful discussions, and critical reading of the manuscript. We thank Steve Siegelbaum for his support and Tobias Bock at Columbia University for providing electrophysiology training to AMP. We thank Gerardo Perera for technical assistance.

This work was supported by Fundação BIAL (Grant 161/10-2010 to I.I.); Fundação para a Ciência e a Tecnologia (PTDC/SAU-NMC/122035/2010 to I.I.); the National Institutes of Health/NINDS (R01NS112485 to I.I.); Consejo Nacional de Ciencia y Tecnología (Grant 254878 to Y.R.C.); and Dirección General de Asuntos del Personal Académico (DGAPA)-PAPIIT, Universidad Nacional Autónoma de México (UNAM) (Grants IN207420 and IN209223 to Y.R.C.).

